# Tonotopically distinct OFF responses arise in the mouse auditory midbrain following sideband suppression

**DOI:** 10.1101/2025.07.02.662630

**Authors:** Patrick D. Parker, Amanda M. Lauer, Dwight E. Bergles

## Abstract

The parsing of sensory information into discrete topographic domains is a fundamental principle of sensory processing. In the auditory cortex, these domains evolve during a stimulus, with the onset and offset of tones evoking distinct spatial patterns of neural activity. However, it is not known where in the auditory system this spatial segregation occurs or how these dynamics are affected by hearing loss. Using widefield single photon neuronal Ca^2+^ imaging in the inferior colliculus (IC) of awake mice, we found that pure tone stimuli elicited both spatially constrained neural activity within isofrequency bands and simultaneous sideband suppression. At cessation of the stimulus, offset responses emerged within the region of sideband suppression, demonstrating that simple stimuli elicit spatiotemporally distinct neural activity patterns to represent the presence of sound and sound termination. Because sound frequency is spatially encoded in the IC, this spatial shift creates a tonotopically distinct offset (tdOFF) response relative to sound onset. Two-photon Ca^2+^ imaging confirmed that tdOFF neuron activity in the sideband region was suppressed during sound and elevated above baseline after stimulus termination, raising the possibility that rebound excitation could contribute to this post-stimulus activation. Loud noise exposure – a common model of hearing loss – abolished both sideband suppression and tdOFF responses. These results show that hearing loss profoundly reshapes the spatiotemporal pattern of sound processing by altering sideband activity. This preferential loss of sideband suppression and tdOFF activation after sound-induced injury in the auditory midbrain may contribute to hyperacusis and tinnitus by promoting neuronal hyperactivity.

**Key Points:**

- Sensory systems encode different features of stimuli by activating distinct neural networks.
- Sound onsets and offsets elicit distinct neural patterns in the auditory cortex, although it is unclear where this separation originates or how it may change with hearing loss.
- Using in vivo widefield Ca^2+^ imaging in awake mice, we find that pure tone stimuli evoke spatiotemporally distinct on-and off patterns of neural activity in the auditory midbrain.
- Neurons active during stimulus offset were suppressed by sound in sideband regions, raising the possibility that rebound excitation contributes to this post-stimulus activation.
- Both sideband suppression and off responses were preferentially abolished following noise-induced hearing loss, raising the possibility that these changes may contribute to hearing loss-related syndromes such as tinnitus and hyperacusis.

## Introduction

Sensory systems separate different elements of sensory information for processing across various brain regions. Within the auditory system, sound frequencies are spatially represented along the cochlear coil, and this tonotopic segregation is maintained throughout auditory centers of the central nervous system (Kandler *et al*., 2009; Kandler, 2018; Babola *et al*., 2018; Kersbergen *et al*., 2022), providing the basis for sound discrimination (Karmakar *et al*., 2017). However, this static view of neural encoding belies their dynamic nature, as populations of neurons adapt to ongoing stimuli to continually encode and process the elements of sound in real time. Neurons in multiple brain regions respond to both the onset (ON) and offset (OFF) of tones. Cortical OFF responses are important for perceiving sound duration and detecting sound termination (Weible *et al*., 2014; Liu *et al*., 2019; Solyga & Barkat, 2021), indicating that the addition of OFF response patterns provides a computational advantage over ON responses or sustained activity alone (Kopp-Scheinpflug *et al*., 2018).

Moreover, individual neurons can have different characteristic frequencies for ON and OFF responses (termed ON/OFF asymmetry) (He, 2002; Scholl *et al*., 2010; Anderson & Linden, 2016; Sollini *et al*., 2018), suggesting that the different temporal periods of the same sound may have distinct tonotopic representations. Indeed, sound ON and OFF pathways are separate in the auditory cortex (AC), have unique tonotopic patterns (Baba *et al*., 2016; Liu *et al*., 2019), and arise from nonoverlapping synaptic inputs (Scholl *et al*., 2010), though it is unclear where in the auditory system these two information streams are spatially segregated and whether impairment of this segregation contributes to hearing disorders.

Subcortical regions that perform early sound processing may establish spatially distinct ON and OFF pathways that are maintained in higher-order regions. OFF responses are present upstream of the cortex in the primary auditory thalamus (medial geniculate nucleus or MGN) (He, 2002; Yu *et al*., 2004; Anderson & Linden, 2016; Solyga & Barkat, 2021) and MGN axons projecting to the primary AC respond to either the ON or OFF period of tones, but seldom both (Liu *et al*., 2019). Whether these separate ON and OFF pathways arise *de novo* in the thalamus or lower brainstem regions is unclear. The inferior colliculus (IC) is the primary source of feedforward input to the MGN and an integrative hub for all brainstem sound processing (Kandler, 2018). Neurons in the central IC are spatially organized into isofrequency lamina, though a subset of neurons show ON/OFF asymmetry, where the characteristic OFF frequency is higher than the ON frequency (Heeringa & van Dijk, 2014; Wong & Borst, 2019), suggesting a systematic shift in the spatial encoding of sound offset toward a lower frequency space. However, it is unknown whether this asymmetry constitutes tonotopically distinct patterns in the IC, but it would suggest that ON and OFF pathways are already separate by the time they exit the brainstem.

Synaptic inhibition may influence the spatial relationship of ON and OFF responses in the IC. Inhibitory GABAergic and glycinergic neurons refine IC sound processing through ‘sideband’ inhibition (Yang *et al*., 1992; Fuzessery & Hall, 1996; Egorova & Ehret, 2008; Pollak *et al*., 2011*b*). Sound-evoked hyperpolarization and suppression of basal firing generates rebound OFF responses in a subset of IC neurons, in part due to hyperpolarization-activated cyclic nucleotide-gated (HCN) channels (Koch & Grothe, 2003; Kasai *et al*., 2012), suggesting sideband inhibition could cluster OFF responding neurons in tonotopic regions adjacent to ON responses. Indeed, OFF responses are more common in IC neurons with sideband inhibition (Akimov *et al*., 2017). Under disease conditions, reduced inhibition is considered a key mechanism of “central gain enhancement” (Auerbach *et al*., 2014), a homeostatic process in the brain that amplifies the remaining peripheral input after hearing loss (Auerbach *et al*., 2014; Palmer & Berger, 2018), which may preferentially alter OFF compared to ON responses (Kopp-Scheinpflug *et al*., 2018). The relationship between IC inhibition and OFF tonotopy is poorly understood, but could contribute to altered neural patterns that underlie hearing loss-related dysfunction, such as poor sound discrimination, persistent phantom sounds (tinnitus), or the overamplification of sound percepts (hyperacusis).

Here, we used single and two-photon (1P and 2P, respectively) imaging of neurons expressing genetically encoded Ca^2+^ sensors in awake mice to define the spatiotemporal dynamics of neural processing of tones in the IC. We found that network activity was highly spatially organized and dynamic, changing its spatial arrangement throughout the stimulus. For many sound frequencies, OFF responses arose from the location of sideband suppression rather than the ON responses, constituting a tonotopically distinct OFF (tdOFF) response pattern. Following loud noise exposure, suprathreshold ON response amplitudes were comparable to pre-noise levels, but sideband suppression and tdOFF responses were abolished, indicating that central gain enhancement has a limited ability to restore these aspects of sound processing. This loss of spatiotemporal inhibition may contribute to neuronal hyperexcitability in hearing disorders resulting from loss of cochlear input.

## Methods

### Neural tracing

To track afferent and efferent projections to the adult mouse IC (postnatal day (P) 56 ± 2), we used the freely available Mouse Brain Connectivity dataset from the Allen Institute for Brain Science (https://connectivity.brain-map.org/) (Oh *et al*., 2014). We used *Cochlear Nuclei (CN)* and *Auditory areas (AUD)* as the *Filter Source Structure* for afferents and efferents, respectively. Experiment #167211503 tracked anterograde projections of parvalbumin (PV) expressing neurons from the CN to the IC using a PV-ires-Cre mouse line. Note that PV is expressed in both excitatory and inhibitory neurons of the CN (Bazwinsky *et al*., 2008; Zhang *et al*., 2022). Experiment #562671482 tracked projections of excitatory neurons from the AC to the outer IC in an Emx1-ires-Cre mouse line.

### Animals

This study was performed in strict accordance with the recommendations provided in the Guide for the Care and Use of Laboratory Animals of the National Institutes of Health. All experiments and procedures were approved by the Johns Hopkins Institutional Care and Use Committee (Protocols #M021M290 & MO24M260). All experiments were performed in adult (8 – 12 weeks old) male and female mice expressing the red-shifted Ca^2+^ indicator jRGECO1a under the Thy1 promoter (Thy1-jRGECO1a mice, line GP8.62) (Dana *et al*., 2018). Mice were maintained on a mixed C57Bl/6NJ;FVB/NJ or C57Bl/6NJ;FVB/NJ;B6.CAST-Cdh23^Ahl+^ background. Genotyping of mice [Thy1-jRGECO1a (Kellner *et al*., 2021) and Cdh23 (Babola *et al*., 2025)] were previously described, and mice contained at least one Ahl allele (Cdh23^Ahl/ahl^ or Cdh23^Ahl/Ahl^) to avoid the early onset of hearing loss associated with Cdh23^ahl/ahl^ in C57Bl/6NJ mice (Babola *et al*., 2025). Mice were housed on a 12-hour light/dark cycle and were provided food ad libitum.

### IC cranial window implant surgeries

Cranial window surgeries to visualize the IC were modified from previous protocols (Goldey *et al*., 2014; Babola *et al*., 2018; Parker *et al*., 2021; Kersbergen *et al*., 2022). Briefly, mice were anesthetized using isoflurane (4 – 5% induction; 1.5% maintenance), the scalp was removed, and a custom metal headbar was affixed to the skull with dental cement (C&B Metabond, Parkell). A circular (3 mm diameter) or square (2 x 2 mm) craniotomy was performed over the left IC using a high-speed dental drill. The dura was retracted, taking care not to damage the transverse sinus or underlying brain. A coverslip [circular: 3 mm diameter, optically glued to a 5 mm diameter (Norland optical adhesive, no. 71); square: 2 x 2 mm glued to a custom stainless steel 17-4PH frame] was placed directly on the IC to stabilize the tissue for imaging, and affixed first with a cyanoacrylate adhesive, then with dental cement. All exposed skull, muscle, and skin margins were sealed with dental cement.

### In vivo Ca^2+^ imaging

Mice were imaged either on the same day of surgery (after > 3 hours of recovery) or within the subsequent 3 days. Mice were head-fixed above a passive disk treadmill (Isett *et al*., 2018) inside a custom-built sound attenuation chamber underneath a Zeiss Axio Zoom.V16 stereo zoom microscope for 1P widefield epifluorescence (Hamamatsu ORCA-Flash4.0 LT digital CMOS camera; Zeiss Illuminator HXP 200C metal halide lamp; 20x – 32x magnification; 10 Hz sampling rate). 2P recordings were performed the same day as the surgery using a Sutter Movable Objective Microscope (Hamamatsu H10770PA-40 SEL and MOD photomultiplier tubes; Coherent Cameleon Discovery laser at 1040 nm excitation, pulse width ≈100 fs, 10 – 20 mW; 620/60 nm emission filters; Nikon 16x/0.8 NA objective and additional 3x digital magnification; field of view 341 x 341 μm, 512 x 512 pixels, 0.67 μm/pixel; 30.05 Hz sampling rate with bilateral scanning).

### Sound stimulation

Sinusoidal amplitude modulated pure tones (10 Hz modulation) were generated with the RPvdsEx software and delivered through a RZ6 Processor and MF-1 free-field speaker (Tucker Davis Technologies) placed 10 cm from the right pinna. Speaker calibration was performed using an ACO Pacific microphone (7017) and preamplifier (4016). Sound stimuli were synchronized with Ca^2+^ recordings using a TTL trigger and consisted of 10 tones at 3 – 32 kHz in ¼ octave intervals and 10 – 16 repeats (mode = 12 repeats) per sound intensity level, cosine-squared gated (5 ms) and played in a random order at 5 s intervals. Unless stated otherwise, the stimulus duration was 1 s (range 0.04 – 3.0 s). Sound intensities were 25 – 105 dB SPL in 20 dB increments.

### Noise exposure

Loud noise exposure was used as a model of induced hearing loss and was performed similar to (Burke *et al*., 2022). Briefly, mice were placed in either a small wire cage or a standard plastic housing cage (30 x 19 x 13 cm) inside a custom-built soundproof chamber (53.5 x 54.5 x 57 cm) while a speaker (Fostex FT28D) suspended above the center of the cage played a narrowband white noise (8–16 kHz) continuously for 2 hours. Mice in the housing cage were allowed to move freely within the cage during the exposure. Sound intensity levels were calibrated using a Larson Davis sound level meter (System 824) and measured 110 – 113 dBA peak level in the center and 103 dBA in the corners. Baseline IC widefield Ca^2+^ recordings were performed before noise exposure and compared to two days after exposure. ABR recordings were performed one day after exposure and confirmed the noise-induced increase in sound-evoked thresholds (Figure 5B). Separate mice were used for IC Ca^2+^ imaging and ABR recordings because the implanted headbar and window required for IC imaging prevented electrode placements for ABR recordings, though mice from both groups were included in each noise trial to ensure they received the same exposure. Control mice for ABR recordings were placed in the noise chamber for two hours under similar conditions, but without the noise stimulus turned on.

### Auditory brainstem response (ABR) recordings

ABR recordings and analysis were performed as previously described in (Kersbergen *et al*., 2022), one day after noise exposure in exposed mice and control, unexposed littermates. Briefly, mice were anesthetized with a mix of ketamine (100 mg/kg) and xylazine (20 mg/kg) delivered by intraperitoneal injection and maintained on a 37°C isothermal heating pad inside a sound attenuation chamber (IAC Acoustics). Platinum needle electrodes (E2, Grass Technologies) were placed subdermal posterior to the left pinna, at the vertex, and in the right leg (ground). Clicks (0.1 ms) or tone pips (5 ms; 2 ms rise time) were presented at 4, 8, 16, 24, and 32 kHz at a rate of 40 Hz with the free-field speaker placed in front of the animal, 10 cm from the vertex of the head. Signals were amplified (Medusa 4Z, Tucker Davis Technologies), band-pass filtered (300 Hz and 3 kHz), digitized (RZ6 Processor, Tucker Davis Technologies), and averaged across 500 stimuli. Evoked thresholds were calculated with custom programs in MATLAB and used linear interpolation to determine the lowest sound intensity with peak-to-peak amplitudes > 2 standard deviations over the background signal.

### Image processing and analysis

Processing and analysis of 1P recordings used custom programs in Fiji and MATLAB. Image files were converted to .tif format and bleach corrected across the entire recording using an exponential fit. Full recordings were divided into individual stimulus trials and normalized as ΔF/F0, where F was an individual frame in the image stack, F0 was the mean fluorescence image from 1 s before stimulus onset, and ΔF = F – F0. Normalized trials were organized by stimulus frequency, aligned to the onset of the tone, and averaged across trials to create a mean image stack of fluorescence changes over time for each frequency (referred to as the trial average).

Response maps for a given sound intensity were from the trial average image stack following a small 3D Gaussian blur (sigma = 1, 1, 0.5 for X, Y, Z), and were the maximum fluorescence change during the tone (referred to as ON maps because the largest amplitude response was always at tone onset) or the 1 s following tone offset (OFF maps). Regions of interest (ROI) to quantify fluorescence responses over time were generated for each stimulus frequency by creating a grand mean of ON maps from 65 – 105 dB SPL trials, normalizing the resulting map to the maximum fluorescence within the IC, and thresholding the map at 0.4 – 0.5 normalized ΔF/F0. The resulting ROIs included both ON and tdOFF areas for a given tone. ON and OFF map contours (Figures 3D and 3F) were similarly generated based on 4% and 7% ΔF/F0 thresholds.

Fluorescence traces were collected as the mean pixel intensity within an ROI from the individual ΔF/F0 stimulus trials, and traces were averaged across trials for quantification. All reported traces are either trial averages from individual animals or grand means of trial averages across animals. In addition, we confirmed ON responses, mid-tone suppression, and tdOFF responses in image stacks and traces from individual trials and the raw recordings. Reported variance is the standard error of the mean. Peak amplitudes of the response were the maximum fluorescence change during the tone and could always be attributed to the ON response. Suppression amplitude was the minimum fluorescence during the tone, and OFF amplitudes were measured from peak suppression (minimum fluorescence) to the maximum fluorescence during the 1 s following tone offset. Rate level functions before and after noise exposure were plotted as the peak amplitude of the ON response for increasing sound intensities (Figure 5D).

We generated kymographs by determining the line perpendicular (running lateral to medial) to the isofrequency bands (running anterior to posterior) in individual animals (see Figure 3C), then averaging all pixels anterior and posterior to the line in the IC, maintaining the lateral to medial spatial information. Spatial profiles of fluorescence intensity across the IC (Figures 6F and 6G) were similar to kymographs and were generated using the same perpendicular line across the ON and OFF response maps.

Image processing and analysis of 2P recordings were the same as 1P, except that ROIs were generated around the visible somas of individual neurons using the average intensity projection over time and in the trial average image stacks. Area under the curve (AUC) was the integral of fluorescence over time [(ΔF/F0)/s] for trial average traces, and the AUC map (Figure 4F, right) was the integral for individual pixels during the tone.

Final figures were designed and generated in Adobe Illustrator.

## Results

### Major divisions of the mouse IC are accessible to *in vivo* optical recording methods

To define the neural activity patterns induced by sound in the auditory midbrain, we first examined the organization of projections within the regions of the IC accessible to *in vivo* 1P and 2P microscopy in adult mice, ~ 300 – 400 µm of the pial surface (Waters, 2020). The central nucleus is the IC segment of the auditory lemniscal pathway, as it combines inputs from multiple brainstem nuclei that process sound information, and it projects to the ventral MGN (Ito *et al*., 2018). In rodents, histochemistry suggests the central nucleus extends nearly to the dorsal surface of the IC (Cant & Benson, 2005; Loftus *et al*., 2008; Lesicko *et al*., 2016), meaning that primary auditory neurons in the IC may be accessed by *in vivo* optical recording methods.

To confirm the anatomy of the mouse IC, we referred to the Allen Institute for Brain Science’s Mouse Connectivity Atlas, a freely accessible database of whole-brain neural tracing experiments (Oh *et al*., 2014). Focal injections of adeno-associated virus (AAV) expressed enhanced green fluorescent protein in neurons of the cochlear nucleus (CN), which is the first brain region to process sound information and a major projection to the central IC (Ryugo & Milinkeviciute, 2023). Reconstructed serial 2P tomography traced projections from the dorsal and ventral CN (DCN and VCN, respectively) to other auditory brain regions, including the contralateral superior olivary complex (SOC), the nucleus of the lateral lemniscus (NLL), and IC (Figure 1A). CN afferents in the IC formed a dense plexus in the central nucleus, extending dorsally to ~ 80 – 100 µm below the pial surface of the IC (Figure 1A, right), consistent with previous functional recordings of tone-evoked responses in putative central IC neurons at this depth in mice (Barnstedt *et al*., 2015).

**Figure 1.**
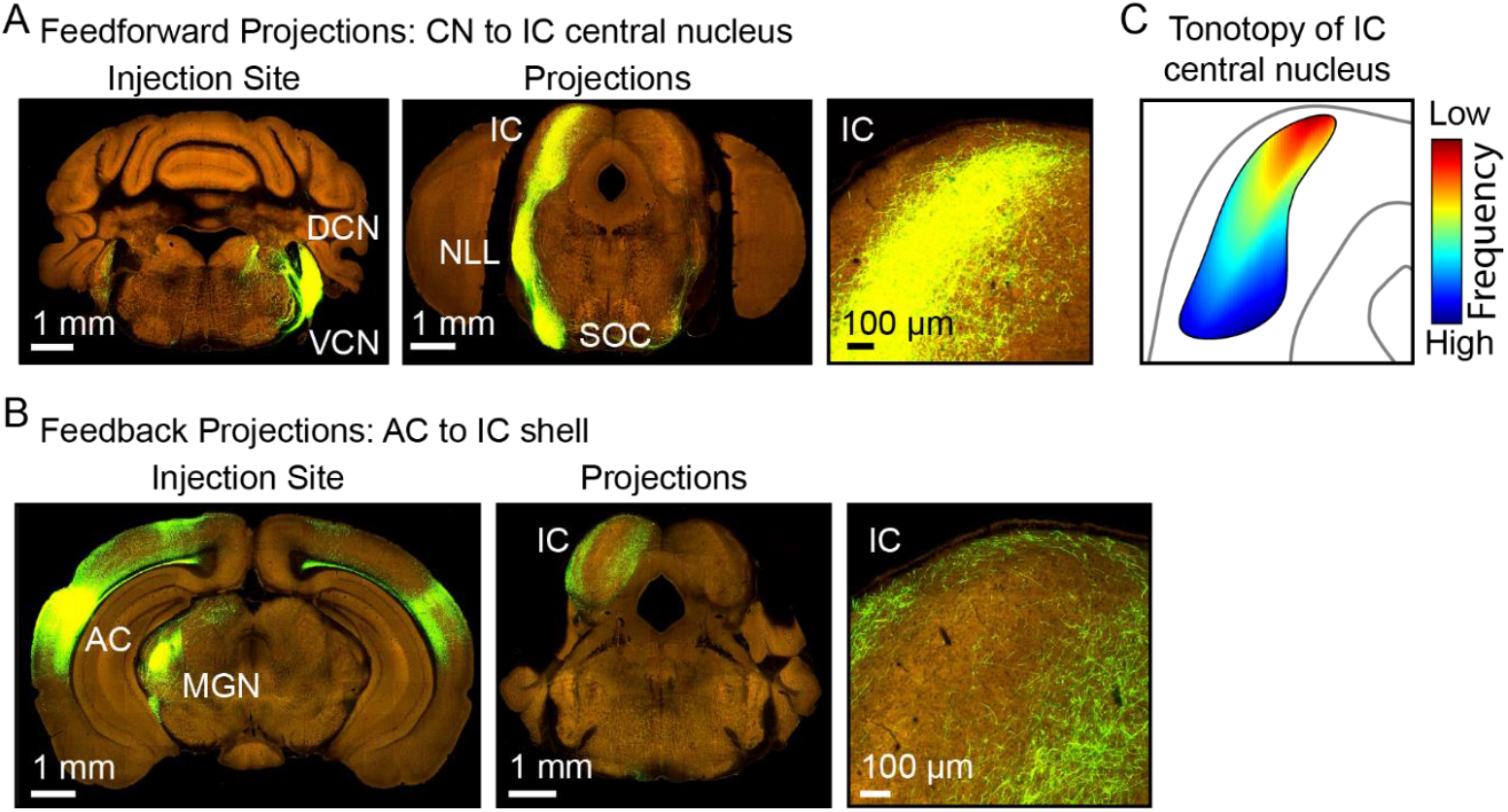
Major divisions of the mouse inferior colliculus are optically accessible. **A**. Feedforward projections from the dorsal and ventral cochlear nucleus (DCN & VCN) project directly to the central nucleus of the IC [Allen Mouse Brain Connectivity Atlas, experiment #167211503; PV-ires-Cre & AAV mediated EGFP expression. Note that PV is expressed in both excitatory and inhibitory neurons of the CN (Bazwinsky *et al*., 2008; Zhang *et al*., 2022)]. **B**. Feedback projections from the auditory cortex (AC) to the lateral and dorsal shell of the IC (experiment #562671482; Emx1-ires-Cre). **C**. Schematic of tonotopic representation in the central IC. NLL = nucleus of the lateral lemniscus; SOC = superior olivary complex; MGN = medial geniculate nucleus of the thalamus.

CN afferents are not restricted to the central IC and may also extend into the lateral cortex and dorsal nucleus regions that form the IC’s outer shell (Ryugo & Milinkeviciute, 2023). This outer shell is distinct from the central IC in that it receives dense corticofugal feedback projections from the AC (Barnstedt *et al*., 2015; Asokan *et al*., 2018). We confirmed the anatomy of the outer shell by tracing AC efferents to the IC and found that they formed a thin band of dense projections limited to < 100 µm below the pial surface (Figure 1B, right). AC efferents provided an outline of the central nucleus and an inverse pattern compared to CN afferents, suggesting that only a thin portion of the outer shell covers the dorsal surface of the central IC nucleus in mice.

The central IC is tonotopically organized in a dorsal-ventral gradient, starting with lower sound frequencies represented in the most superficial regions and progressively higher frequencies represented at greater depths (Portfors *et al*., 2011) (Figure 1C). The innervation patterns of CN afferents and AC efferents suggest that neurons processing low-frequency sound information along the primary lemniscal pathway in the IC reside starting ~ 100 µm or less below the pial surface, an accessible depth for *in vivo* fluorescence microscopy (Barnstedt *et al*., 2015; Wong & Borst, 2019; Waters, 2020).

### Widefield Ca^2+^ imaging reveals spatiotemporal patterns of sound processing in the IC

To understand the spatiotemporal characteristics of neural sound processing in the IC, we used 1P widefield epifluorescence imaging to visualize sound-evoked Ca^2+^ transients in awake Thy1-jRGEGO1a mice (Dana *et al*., 2018) (Figure 2A). Amplitude-modulated tones (3 – 32 kHz; 25 – 105 dB SPL; 10 Hz modulation) presented to the contralateral ear elicited an increase in neural Ca^2+^ in spatially discrete domains along presumed isofrequency laminae (Figure 2B). The lowest frequencies (3 – 4 kHz) evoked 1 – 2 parallel bands of neural activity obliquely oriented to the rostro-caudal axis. Higher sound frequencies (~ 5 – 16 kHz) evoked two spatially distinct bands that were increasingly separated from one another as we increased the stimulus frequency, similar to previous functional recordings (Babola *et al*., 2018; Wong & Borst, 2019; Kersbergen *et al*., 2022), consistent with the bifurcation of afferents from the brainstem (Malmierca *et al*., 1995; Ryugo & Milinkeviciute, 2023). Merging peak responses for all sound frequencies revealed the nested pattern of tonotopy along the dorsal surface of the IC (Figure 2B, *lower left*), highlighting the spatial specificity of neural response patterns visible with widefield Ca^2+^ imaging. Above ~ 16 kHz, the peak response remained in the same spatial location, but the amplitudes gradually decreased with increased sound frequency (Figures 2B – 2D), likely reflecting the fact that neurons at these characteristic frequencies reside increasingly deeper in the IC along the same vertical axis, and that their responses and susceptible to greater attenuation due to scattering by the overlying tissue.

**Figure 2.**
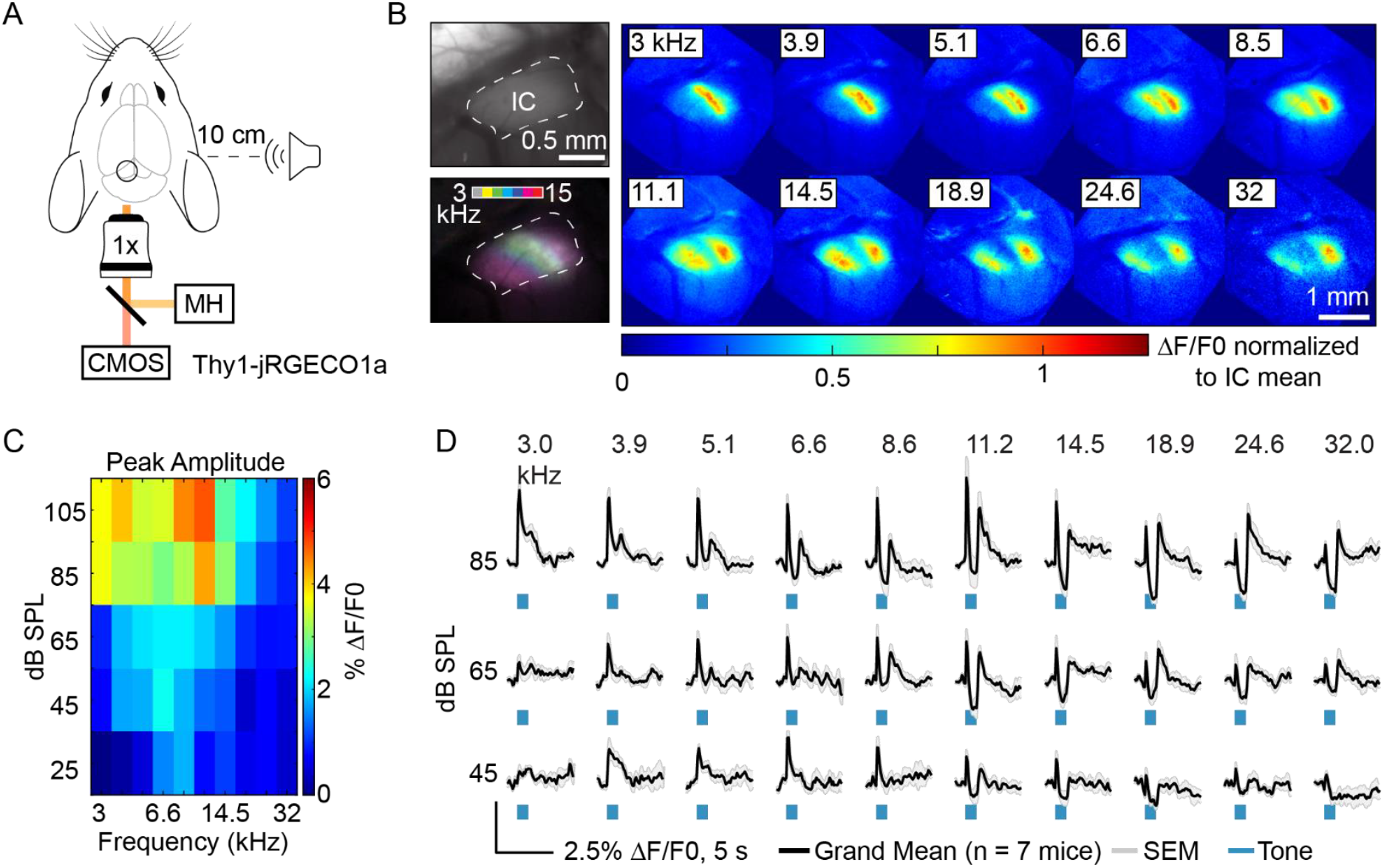
Tonotopic organization of sound-evoked neural activity in the IC using widefield neural Ca^2+^ imaging. **A**. Tone-evoked neural activity monitored in the IC using single photon, widefield Ca^2+^ imaging in awake adult mice (Thy1-jRGECO1a). MH = metal halide lamp; CMOS = complementary metal oxide semiconductor. **B**. Evoked changes in neural Ca^2+^ fluorescence in the IC to individual tones during sound presentation (right; average 65 – 105 dB SPL, 1 s stimulus duration) in a single mouse. Pixel intensity normalized to the average across the entire IC for a given tone. A merged image with pseudocolored responses (lower left) shows the tonotopic organization visualized from the dorsal surface of the IC. Upper left: raw fluorescence image. **C**. Frequency response area of the peak amplitude of evoked changes in neural Ca^2+^ fluorescence during tone presentation (i.e., excluding OFF responses; grand mean of n = 7 mice). **D**. Fluorescence intensity changes over time demonstrate dynamic responses in the IC in a frequency-and sound intensity-dependent manner. Grand mean +/-SEM (n = 7 mice).

Widefield Ca^2+^ imaging recorded evoked neural responses at the lowest sound intensities tested (25 dB SPL) for a subset of frequencies centered at 6 – 8 kHz (Figure 2C). This likely reflects a combination of the mouse’s biological sensitivity to these frequencies and the proximity of these neurons to the dorsal surface (Ralls, 1967; Portfors *et al*., 2011), making them more amenable to optical recording. Increasing sound intensities monotonically increased the peak amplitude of the response for a given frequency (Table 1), and the range of frequencies that evoked a detectable response, creating a “V”-shaped frequency response area (Figure 2C).

**Table 1.**
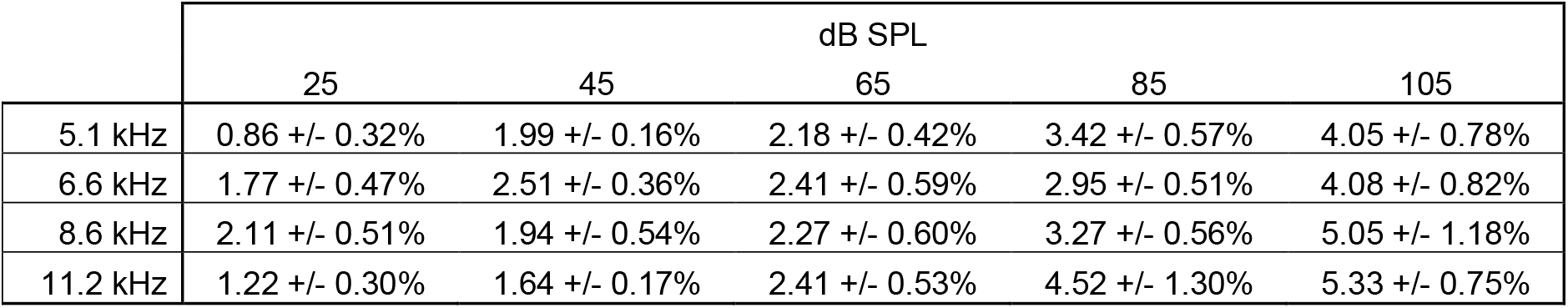
Peak Ca^2+^ response amplitudes for a set of tones with increasing sound intensity. Mean ± standard error of the mean (SEM) as % ΔF/F0.

Spatial maps of the peak responses present a static representation of neural activity during sound processing; however, these patterns evolved with time during and after sound presentation. Across all sound intensities and frequencies, the peak amplitude of the neural response occurred immediately following tone onset (ON) (Figure 2D). During a 1 s duration tone (45 – 85 dB SPL), neural responses quickly adapted and Ca^2+^ fluorescence decreased immediately following the peak. At frequencies greater than 6 kHz, sound-evoked fluorescence dropped below pre-stimulus levels, consistent with sound-induced suppression of basal neural activity (Figure 2D; see also Figures 3 & 4) (Ali & Kwan, 2019). Importantly, tones did not evoke any increase or decrease in fluorescence intensity in control mice lacking any functional indicator in our recording conditions, suggesting the mid-tone suppression of Ca ^2+^ fluorescence was unlikely due to a hemodynamic or movement artifact, but due to the known inhibition of basal firing rates in some IC neurons to tones outside of their characteristic frequency (Kasai *et al*., 2012; Heeringa & van Dijk, 2014; Wong & Borst, 2019). Most tones elicited an OFF response, defined as an increase in neural fluorescence following the cessation of the stimulus. OFF amplitudes were relatively small compared to ON at low stimulus frequencies (e.g., 3 & 3.9 kHz) (Figures 2D & 3G). However, at higher frequencies OFF response amplitudes were comparable to or even exceeded ON responses, measured from either the pre-stimulus fluorescence level or from the point of maximum suppression (e.g., 24.6 & 32.0 kHz).

These results indicate that widefield Ca^2+^ imaging is able to resolve the tonotopic organization and dynamic temporal aspects of neural response patterns during sound processing in the IC, including ON and OFF responses, adaptation, and mid-tone suppression.

### OFF responses are tonotopically distinct from ON responses

To ensure OFF responses were time-locked to the end of the stimulus and to understand the time course of suppression, we varied the duration of the tones (40 ms, 400 ms, 1 s, and 3 s). Temporally distinct OFF responses followed stimuli lasting ≥ 400 ms (Figure 3B), and the rise in fluorescence of the OFF response always started in the recording frame immediately following the cessation of the stimulus (< 100 ms), regardless of the stimulus duration. This confirmed that neurons responded to the withdrawal of sound rather than a delayed response to the onset of the tone. At stimulus durations ≤ 400 ms, it was difficult to distinguish neural adaptation from inhibition, though 1 s and 3 s duration tones consistently evoked a distinct period of neural suppression. During the longest duration tones (3 s), the response reached a maximum suppression, then gradually increased fluorescence toward baseline during the tone (Figure 3B, bottom right), suggesting the level of neural inhibition peaked within the first 1 s of the stimulus and adapted during prolonged stimulation. Thus, both mid-tone suppression and OFF responses depended on stimulus duration. We chose a 1 s tone for the remainder of this study, as it produced peak suppression and a prominent OFF response at most frequencies.

**Figure 3.**
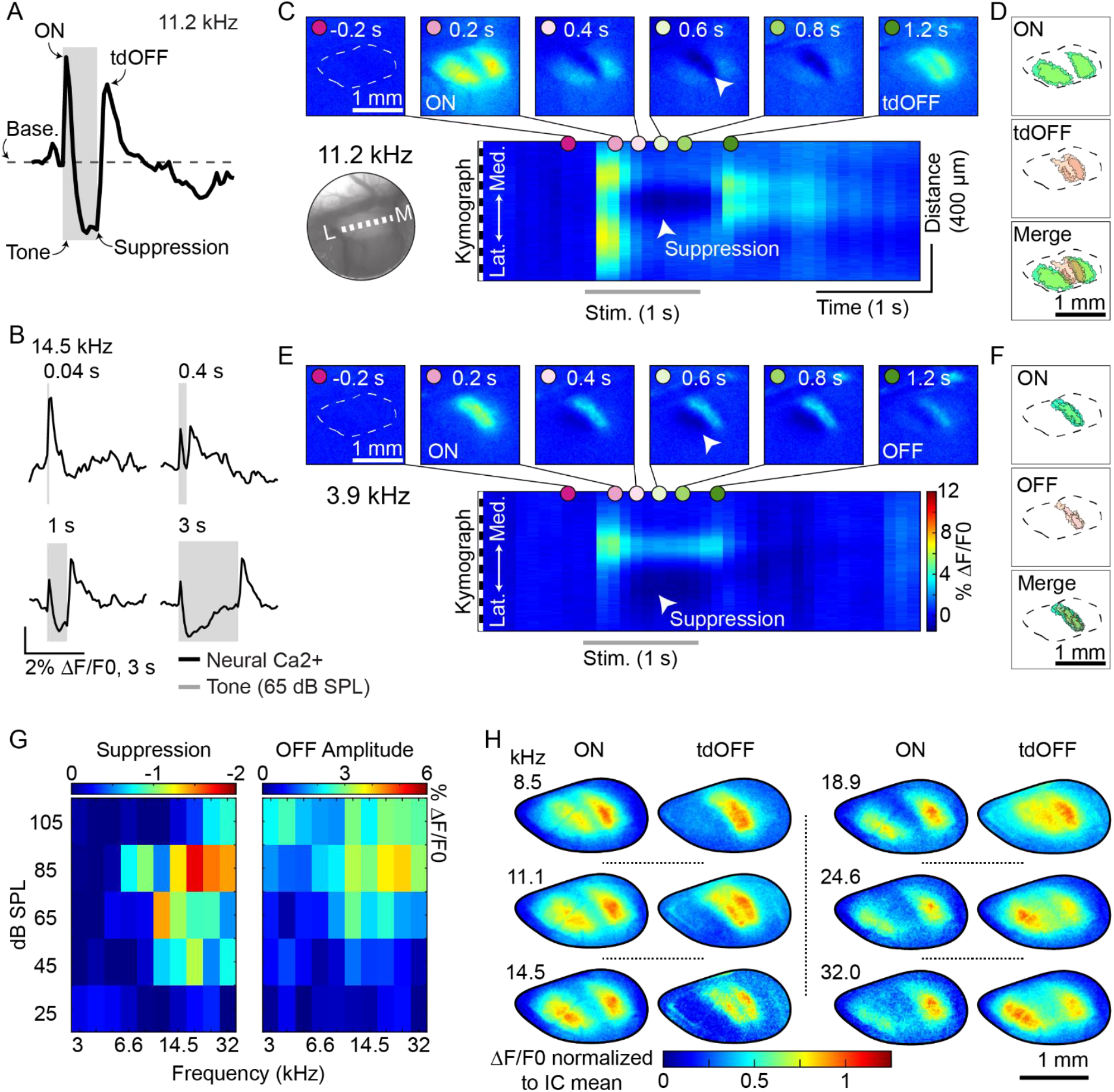
Tonotopically distinct OFF responses (tdOFF) spatially correlate with evoked sideband suppression. **A**. Anatomy of an evoked response, including ON, suppression, and tdOFF responses. Trial averages from the same animal and frequency as C. Base. = baseline fluorescence level. **B**. tdOFF responses based on tone duration (from a single animal). **C**. Spatiotemporal dynamics of neural activity in the IC for a single tone (11.2 kHz) with tdOFF response. Top: individual imaging frames aligned to the start of the stimulus (0 s; tone offset = 1 s). Arrowhead points to tone-evoked sideband suppression of neural Ca^2+^ fluorescence. Bottom: kymograph illustrating the lateral/medial positioning (vertical axis) of neural activity over time (horizontal axis). Note the spatial overlap of suppression and tdOFF, but distinction from ON. Color scale bar as in E. Averaged across trials; 85 dB SPL. **D**. Maps of ON and tdOFF responses illustrate their distinct spatial patterns. Contour lines based pseudocolored maps for approximately 4% and 7% ΔF/F0. Same stimulus as C. (continued next page). **E & F**. Similar to C & D, but for a frequency (3.9 kHz) lacking tdOFF. Note that a small amplitude OFF response spatially overlaps with the ON response at this frequency. **G**. Frequency response areas of peak suppression and tdOFF amplitudes. Grand mean of the same n = 7 mice as Figure 2C. **H**. Comparing the spatial patterns of ON and tdOFF responses over a range of frequencies.

Mid-tone suppression and OFF responses were spatially correlated, with OFF responses arising from the same region as evoked suppression. Tones ≥ 6 kHz evoked an ON and sustained response consisting of two bands of neural activity, as well as an area of suppression located between the two bands – consistent with evoked sideband inhibition in a region tonotopically aligned with a lower sound frequency than the ON response (Figure 3C; an arrowhead marks the area of suppression). To help resolve this spatial relationship over time, we generated kymographs by averaging the anterior/posterior pixel intensities of the IC but maintained the medial/lateral aspects (Figure 3C, *bottom*). At the end of the stimulus, OFF responses had more spatial overlap with the region of evoked suppression than the ON responses (Figure 3C & 3D). We termed this pattern “tdOFF,” as it was tonotopically distinct from the ON and sustained response patterns for the same tone, and to distinguish tdOFF from OFF responses that tonotopically overlap with ON responses. At these stimulus frequencies, the area of suppression and tdOFF were always located in a tonotopic region corresponding to a lower frequency than the ON response (Figure 3H), indicating a systematic spatial shift in neural patterns for the presence vs. termination of sound.

In contrast, the lowest sound frequencies tested (3 – 4 kHz) evoked very little suppression and only a small amplitude OFF response at the highest sound intensities (≥ 85 dB SPL) (Figure 3G). Evoked suppression arose in an adjacent sideband in a tonotopically higher region (rather than lower), and OFF responses were spatially aligned with ON responses (Figure 3E & 3F), indicating tdOFF did not occur at these lowest sound frequencies. This analysis reveals that for most frequencies, sound onset and offset evoke tonotopically distinct patterns of neurons in the IC, and that tdOFF responses correlate with sideband suppression, rather than ON responses. However, widefield imaging does not provide cellular resolution, limiting our interpretation of the spatial relationship of neural response types, as well as the relationship of sound-evoked suppression and tdOFF responses.

### tdOFF-responding neurons are clustered in sideband regions and inhibited by tones

To define the spatiotemporal relationship between evoked suppression and tdOFF responses in individual neurons, we used 2P Ca^2+^ imaging in the IC while presenting tones to awake Thy1-jRGECO1a mice (Figure 4A & 4B). We confirmed that IC neurons that responded to a given tone clustered together in presumed isofrequency laminae (Figure 4C). Ca^2+^ fluorescence in the surrounding neuropil also increased and was spatially restricted, likely due to the coordinated activity of ascending axons and local dendrites. At an imaging depth of ~ 237 µm, a 6.9 kHz tone produced the largest amplitude response and the greatest number of responding neurons. For lower frequency tones, we found a larger number of responding neurons at shallower imaging depths, consistent with previous 2P Ca^2+^ recordings in mice (Barnstedt *et al*., 2015) and supporting the conclusion that the lowest frequency regions of the central IC reside within a few hundred microns of the pial surface (see Figure 1).

**Figure 4.**
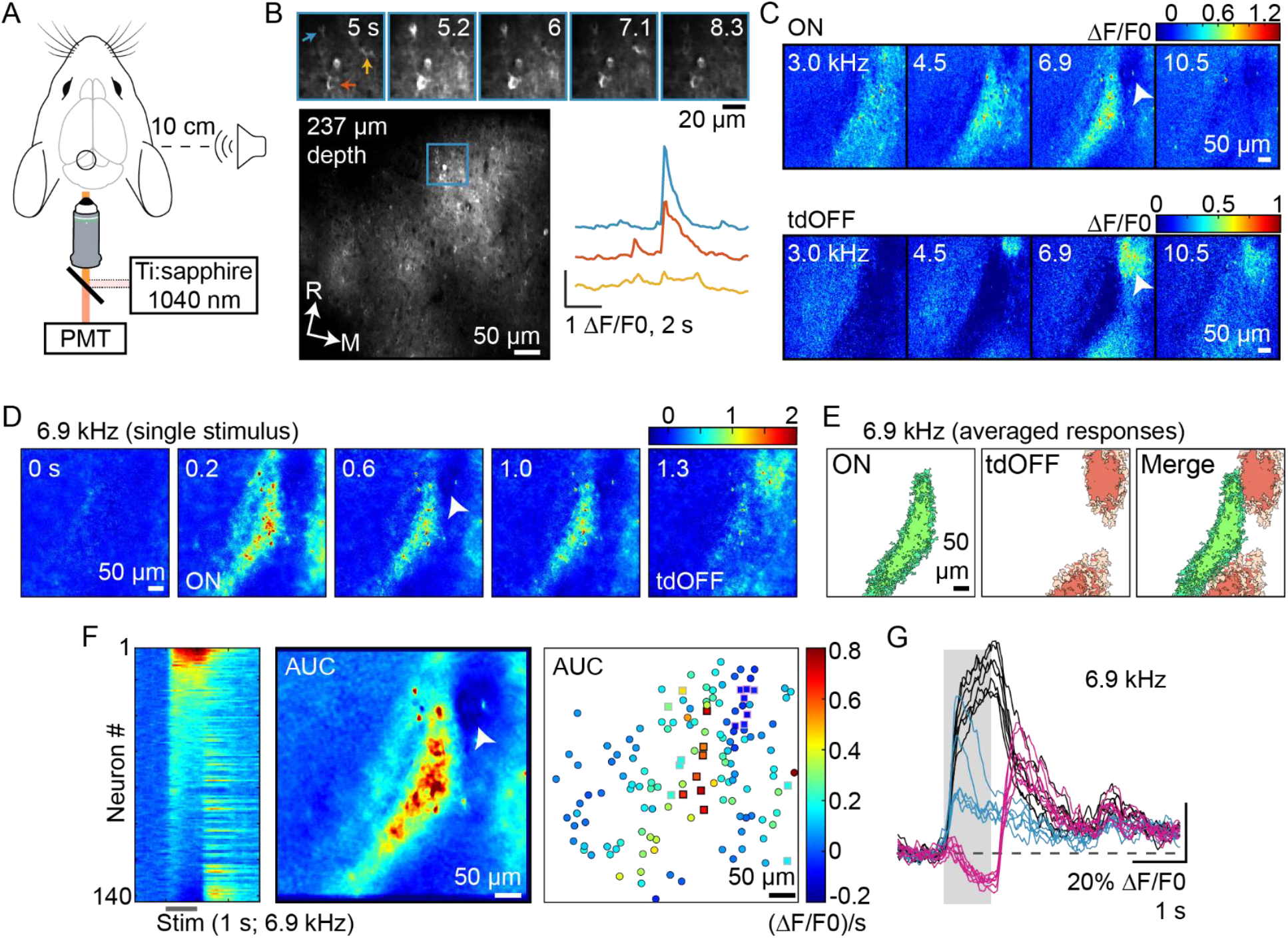
tdOFF responding neurons are suppressed by sound in sideband regions. **A**. Tone-evoked neural activity monitored in the IC using 2P Ca2+ imaging in awake adult mice (Thy1-jRGECO1a). **B**. Averaged intensity projection of all neurons in the field of view (FOV) (bottom left), as well as a series of recording frames (top) and fluorescence traces (bottom right) illustrating Ca^2+^ activity in three individual neurons. R = rostral. M = medial. Time stamp relative to the start of the traces. **C**. Averaged ON and tdOFF response maps for different frequencies in the same FOV as B (85 dB SPL, 1 s stimulus duration). A 6.9 kHz tone evoked the largest amplitude response and recruited the most neuron. Arrowhead marks the location of evoked suppression and subsequent tdOFF. **D**. Individual recording frames illustrate the spatiotemporal dynamics of neural activity during a single stimulus. Time stamp relative to tone onset (1 s duration). **E**. Contour maps of ON and tdOFF responses to 6.9 kHz tone illustrate their distinct spatial patterns. Based on averaged maps in C. **F**. Types of neural responses spatially cluster. Left: raster plot of neural fluorescence changes over time for all neurons in the FOV, ordered by their integral (‘area under the curve’; AUC) during the 6.9 kHz tone. Center: AUC response map. Right: spatial positions of neurons (circles and squares) and their AUC (color). Square neurons are quantified in G. **G**. Fluorescence intensity traces for select neurons based on their spatial positioning (squares in F) and response type. Black: neurons with a continued increase in fluorescence during the tone (black) from the ON isofrequency band. Blue: adapting neurons from the edge of the ON band (grey squares in F). Pink: tdOFF neurons from the sideband region with peak suppression.

The 6.9 kHz tone suppressed Ca^2+^ fluorescence in a cluster of neural somas and neuropil in a sideband region adjacent to the responding lamina (arrowhead in Figures 4C & 4D), which also became the location of the tdOFF response at tone offset (Figure 4C – 4E). To better understand how spatial positioning influenced neural response patterns to the tone, we sorted all the active neurons in the field of view based on the integral of their somal Ca^2+^ fluorescence during the tone (Figure 4F, *left*). OFF responses were visible in a subset of neurons, including those inhibited by the tone and some neurons with ON-adapting responses, consistent with previous reports (Xie *et al*., 2007; Kasai *et al*., 2012; Wong & Borst, 2019). Neurons with the highest integral resided together in the responding laminae (Figure 4F, *right*; square ROIs with black outline) and continuously increased their Ca^2+^ fluorescence for the duration of the tone (Figure 4G, *black*), while neurons at the edge of the lamina showed more adaptive responses (grey ROI outlines & blue traces), suggesting that spatial positioning influences the response characteristics of a neuron to a given sound frequency. Neurons with the lowest or negative integrals were spatially clustered together in the sideband region of maximum suppression and contained a prominent tdOFF response (pink outline and traces).

Two-photon imaging confirmed that tone-evoked suppression precedes tdOFF responses in individual neurons and demonstrated, for the first time, that these neurons are clustered in a tonotopic region distinct from ON responses in the IC.

### Hearing loss disrupts evoked suppression and tdOFF responses more than ON responses

With hearing loss, a compensatory reduction in inhibitory function and machinery along the auditory pathway is thought to help restore suprathreshold responses to normal hearing levels (Auerbach *et al*., 2014; Palmer & Berger, 2018). Given the spatiotemporal relationship of evoked suppression and tdOFF responses (Figures 3 & 4), we explored how hearing loss might influence tdOFF responses. We chose a model of loud noise-induced hearing loss (8 – 16 kHz at 110 dB SPL for 2 hours) (Figure 5A), which causes a permanent increase in sound-evoked thresholds (Ohlemiller *et al*., 2000) and a reduction in inner hair cell synapses in young adult mice within 24 hours (Wu *et al*., 2024). Noise exposure diminished ABR waveform amplitudes and significantly increased evoked thresholds compared to control (unexposed) littermates one day after exposure (two-way ANOVA for the effect of exposure condition and stimulus frequency, including click, on ABR thresholds: exposure *F* (1, 49) = 101.84, p = 1.49e^-13^, *η*^2^ = 0.61; stimulus frequency *F* (5, 49) = 2.11, p = 0.08, *η*^2^ = 0.06) (Figure 5B), demonstrating a significant impairment in the ability of the inner ear to transduce or convey external sounds to the central nervous system, independent of the sound frequencies tested. In a separate set of mice, we used widefield Ca^2+^ imaging to compare sound-evoked IC responses before and after noise exposure in the same animal. Evoked Ca^2+^ thresholds in the IC were affected similarly to ABRs two days after exposure, requiring sound intensities ≥ 85 dB SPL to elicit responses for many frequencies (two-way ANOVA on evoked Ca^2+^ thresholds: exposure *F* (1, 89) = 342.11, p = 3.05e^-32^, η^2^ = 0.78; stimulus frequency *F* (9, 89) = 1.09, p = 0.38, η^2^ = 0.02) (Figures 5C & 5D). Despite the elevated thresholds, the amplitude of suprathreshold Ca^2+^ responses either matched pre-exposure levels (3.9 – 14.5 kHz) or were potentiated (18.9 – 32 kHz) (Figure 5D), consistent with the amplification of neural responses to match pre-exposure levels and a hallmark of central gain enhancement (Auerbach *et al*., 2014; Chambers *et al*., 2016; Asokan *et al*., 2018).

**Figure 5.**
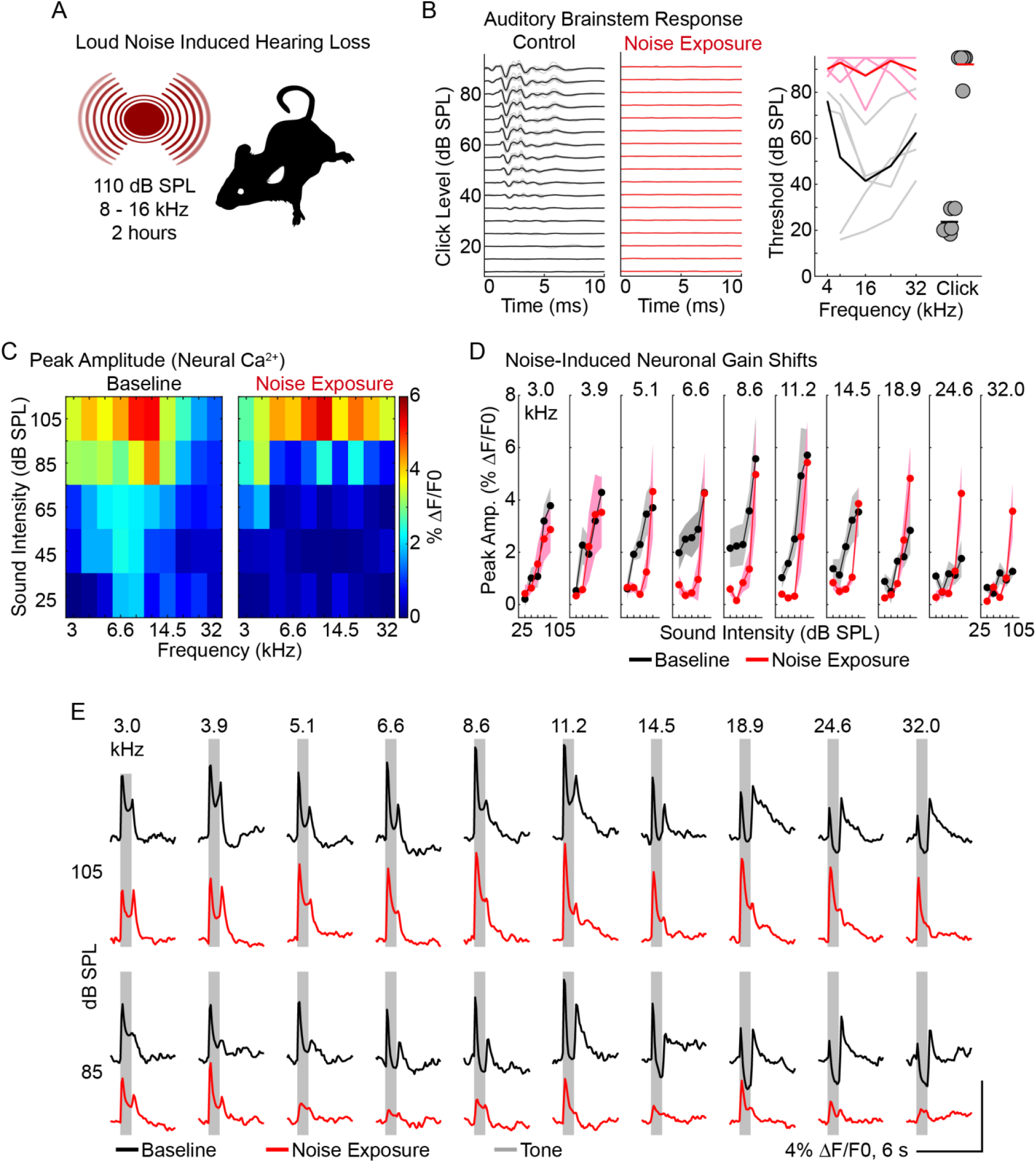
Loud noise-induced hearing loss alters the temporal characteristics of evoked responses. **A**. Loud noise exposure was used as a model of sound-induced hearing loss. **B**. Auditory brainstem response (ABR) electrical recordings demonstrate the elevated sound-evoked thresholds indicative of hearing loss following noise exposure compared to non-exposed (control) littermates. Left: grand mean ABR traces (dark lines) and standard error (gray lines) for both groups. Right: evoked thresholds to a broadband click and individual tones. n = 5 mice. Recorded one day after noise exposure. (continued next page). **C**. Frequency response area of the peak amplitude of evoked changes in neural Ca^2+^ fluorescence during tone presentation in the same mice measured before (Baseline) and two days after noise exposure (grand mean of n = 5 mice). **D**. Rate-level functions measured as the amplitude of peak fluorescence at increasing sound intensities. The increased slope for many frequencies and potentiated response ≥ 18.9 kHz indicate enhanced neural gain following noise exposure. Grand means and standard error. **E**. Grand mean fluorescence intensity traces at suprathreshold sound intensity levels for both groups. Note the lack of evoked suppression and tdOFF at most frequencies after noise exposure.

Loud noise exposure caused a striking alteration in the temporal response patterns of sound processing in the IC. Similar to baseline recordings, ON responses accounted for the peak amplitude of evoked fluorescence after noise exposure, but evoked Ca^2+^ fluorescence never dropped below pre-stimulus levels for any frequency, consistent with a significant reduction in evoked inhibition (Figures 5E & 6A). Instead of suppression, neurons adapted to a higher amplitude steady state following noise exposure (Figures 6C and 6D). OFF response amplitudes (including OFF and tdOFF) were diminished or absent at most sound frequencies (Figures 5E & 6B), despite comparable ON amplitudes to baseline (Figure 5D & 5E), suggesting evoked suppression and OFF responses were more affected by noise exposure than ON responses.

**Figure 6.**
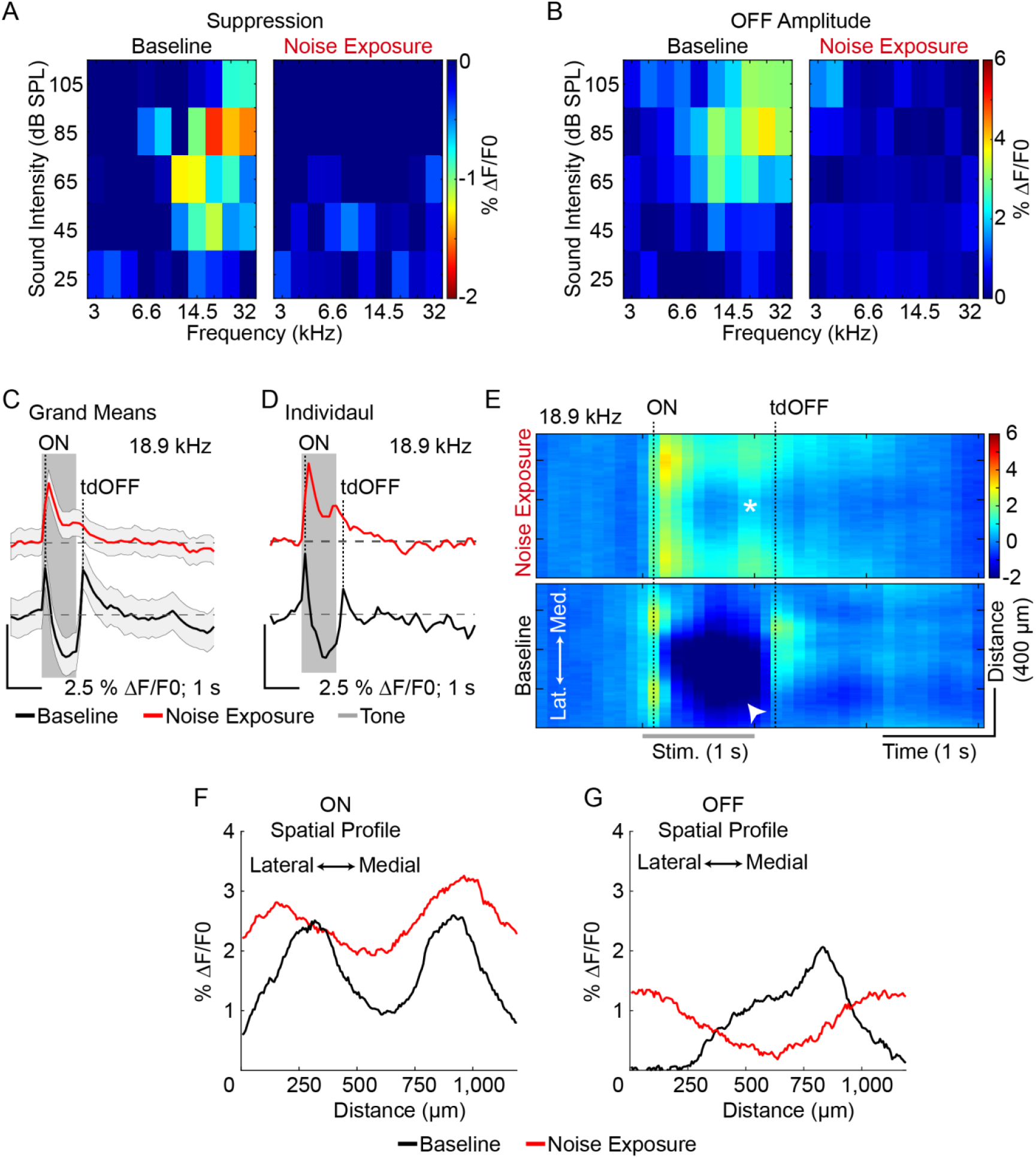
Loud noise-induced hearing loss abolishes sideband suppression and tdOFF responses. **A & B**. Frequency response areas of peak suppression (A) and OFF amplitudes (B) before and after noise exposure in the same mice. Note the lack of both response features after noise exposure. Grand mean of the same n = 5 mice as Figure 5. **C & D**. Fluorescence intensity traces for a given frequency (18.9 kHz) highlight the temporal change in neural responses due to the loss of evoked suppression and tdOFF after noise exposure. C = grand mean and standard error. D = trial average for an individual mouse. Dashed lines = basal fluorescence level before the tone. **E**. Kymographs highlight the spatial change in neural response patterns after noise exposure. Arrowhead = evoked neural suppression. * = expansion of the response area during the tone. Note that after noise exposure, neural fluorescence persists after tone offset at the ON location. **F & G**. Noise exposure alters the neural spatial profile – a measure of fluorescence intensity across the IC from lateral to medial. F = spatial profile at the peak response during the tone (ON). G = spatial profile of the max fluorescence following tone offset.

Noise exposure abolished tdOFF responses and reshaped the spatial patterns of sound processing. With the loss of evoked suppression after noise exposure, response areas persisted and, in some instances, expanded in size during the tone (asterisk in Figure 6E, *top*), a phenotype not seen in recordings prior to noise exposure. tdOFF responses were also absent after noise exposure, and neural Ca^2+^ activity remained elevated at the location of ON responses (Figures 6E & 6G), severely altering neural patterns marking the end of the sound stimulus.

These results indicate that the spatiotemporal dynamics of sound-evoked neural activity are severely disrupted in the IC after damaging noise exposure. While suprathreshold ON response amplitudes recovered to baseline levels, noise exposure abolished evoked suppression and the associated tdOFF responses, providing further supporting evidence for a relationship between inhibition and tdOFF responses, and suggesting that central gain enhancement may prioritize ON responses over tdOFF after hearing loss.

## Discussion

OFF responses are present across sensory systems throughout the mammalian nervous system and are important to discriminate features and distinguish the end of a stimulus. Within the auditory system, OFF responses comprise a distinct pattern of neurons and maintain a larger spatial area than ON responses in the primary, secondary, and associated AC (Baba *et al*., 2016; Sollini *et al*., 2018; Liu *et al*., 2019). AC cortical neurons receive nonoverlapping ON and OFF synaptic inputs (Scholl *et al*., 2010), and MGN afferents – the primary source of sound information to the cortex – are biased toward either ON or OFF responses (Liu *et al*., 2019), suggesting separate, but parallel pathways for processing sound onset and offset could arise upstream in the brainstem. We utilized *in vivo* 1P and 2P fluorescent Ca^2+^ imaging in mice to understand the spatiotemporal dynamics of sound processing in the IC during normal and impaired hearing. We show that the central nucleus of the IC is accessible to *in vivo* optical recording methods and that, with normal hearing, tones evoke a tonotopically discreate pattern of neural activity, followed by a tonotopically distinct offset response pattern (tdOFF), demonstrating for that these pathways spatially separate in the midbrain before ascending to the thalamus and AC. Widefield imaging showed that tdOFF responses spatially overlapped with sideband suppression, and 2P imaging confirmed that tdOFF neurons were suppressed by tones, suggesting a potential link between inhibition and tdOFF responses, as well as providing a focus for future studies to determine the mechanism(s) that generate tdOFF patterns. Noise-induced hearing loss altered the landscape of sound processing in the IC, and tdOFF responses appeared to be more sensitive than ON responses to the decrease in peripheral input, as noise exposure abolished side-band suppression and tdOFF responses, but maintained ON response amplitudes comparable to baseline recordings at suprathreshold sound intensities. It remains to be seen whether impaired tdOFF responses contribute to known perceptual issues associated with aging and hearing loss, such as speech discrimination, hyperacusis, or tinnitus.

### Inhibition and IC OFF responses

The IC receives synaptic inhibition from local GABAergic neurons, as well as glycinergic and GABAergic projection neurons from the nucleus of the lateral lemniscus, superior paraolivary nucleus (SPN), and the contralateral IC (Yang *et al*., 1992; Fuzessery & Hall, 1996; Palombi & Caspary, 1996; Xie *et al*., 2007; Egorova & Ehret, 2008; Pollak *et al*., 2011*a*, 2011*b*). The spatial correlation between sideband suppression and tdOFF responses suggests that evoked inhibition may spatially cluster tdOFF-responding neurons. Indeed, the IC contains the machinery for inhibition-mediated neural excitation: neurons express multiple HCN channels (HCN1, HCN2, and HCN4), and a subset of excitatory neurons contain fast activating, large amplitude hyperpolarizing-activated currents (Ih) sufficient to generate rebound action potentials *in vitro* (Koch & Grothe, 2003; Naumov *et al*., 2019). *In vivo*, tone-induced inhibition and suppression of baseline firing induces rebound action potentials without excitatory synaptic input in a subset of IC neurons (Kasai *et al*., 2012), suggesting inhibition alone is sufficient to induce OFF responses. Indeed, OFF responses are more common in IC neurons with sideband inhibition (Akimov *et al*., 2017), and the magnitude of the OFF response (i.e., number of action potentials) increases with the duration of sound (Kasai *et al*., 2012). We found that tones suppressed tdOFF neurons in sideband areas, suggesting that synaptic inhibition may summate in this region to facilitate rebound responses (Sanchez *et al*., 2008), spatially clustering tdOFF responding neurons in tonotopic regions adjacent to ON isofrequency lamina.

However, not all IC OFF responses require inhibition or hyperpolarization, and OFF responses derive from multiple mechanisms. Some OFF responses are unaffected by local pharmacological blockage of ionotropic GABA and glycine receptors or are due to synaptic excitation timed to sound termination (Vater *et al*., 1992; Kasai *et al*., 2012). Our 2P recordings similarly found neurons containing OFF responses without tone-evoked suppression, though these neurons were spatially separate from tone-suppressed tdOFF responses. In these instances, the amplitude of OFF responses depended on a neuron’s proximity to the area of maximal suppression, similar to the strength of ON responses depending on the spatial proximity to the evoked isofrequency lamina. Could the spatial location of a neuron and its proximity to the evoked isofrequency lamina determine the mechanism(s) of OFF responses and their dependence on inhibition? Both synaptic excitation arriving at the end of a tone and inhibitory mechanisms can, by themselves, generate OFF responses, though combining synaptic and rebound excitation in the same neurons could potentiate OFF responses (Kopp-Scheinpflug *et al*., 2018). In this scenario, inhibition is required to both generate the largest amplitude OFF responses and coordinate them in a tonotopically distinct region from ON responses.

While tdOFF responses correlated with evoked suppression in both widefield and 2P recordings, we did not test the role of synaptic inhibition or determine the underlying mechanism(s). Inhibition is crucial for sound processing in the IC (Pollak *et al*., 2011*b*), and determining the influence on tdOFF responses will inform how inhibition aids in encoding the end of sounds.

### OFF responses in the auditory brainstem

Brainstem regions that process sound information upstream of the IC also contain OFF responses, including the cochlear nucleus (CN) (Suga, 1964; Young & Brownell, 1976; Ding *et al*., 1999), superior olivary complex (Covey *et al*., 1991; Kuwada & Batra, 1999; Kopp-Scheinpflug *et al*., 2011), and nuclei of the lateral lemniscus (Covey & Casseday, 1991), all of which are likely to influence IC processing. Both the DCN and VCN provide glutamatergic projections to the IC, and CN OFF responses are a potential source of synaptic excitation for tdOFF response in the IC (Vater *et al*., 1992; Kasai *et al*., 2012; Kopp-Scheinpflug *et al*., 2018). However, afferents from the CN are believed to project to the same tonotopic region of the IC (Ryugo & Milinkeviciute, 2023), and we find tdOFF responses in sideband regions, meaning CN OFF responses may have a greater influence on IC OFF responses spatially aligned with ON responses, though this remains to be tested. The SPN contains the most well-characterized OFF response in the auditory system, where sounds evoke glycinergic synaptic inhibition from the medial nucleus of the trapezoid body and suppress action potential firing in the SPN (Kopp-Scheinpflug *et al*., 2011). The termination of sound and inhibition results in rebound OFF responses due to large Ih currents and T-type Ca^2+^ conductance. SPN neurons are GABAergic and OFF responses provide synaptic inhibition to the IC at sound offset (Felix *et al*., 2015), meaning they likely influence continued neural activity following sound offset rather than generate tdOFF responses in the IC. It remains to be determined whether the tdOFF responses arise from upstream brainstem inputs or de novo in the IC.

### Potential consequences of tdOFF responses

The IC is an integrative hub of sound processing pathways from the brainstem, and any OFF response from the brainstem would have to pass through the IC before reaching the thalamus and cortex (though a few brainstem projections circumvent the IC (Malmierca *et al*., 2002)). Transgenic mouse lines based on the Thy1 promoter express in glutamatergic neurons in most brain regions (Dana *et al*., 2018), including the IC (Wong & Borst, 2019). Projections from the IC reach the thalamic MGN, but the IC has no apparent innervation of the inhibitory reticular nucleus (Ledoux *et al*., 1987; Cornwall *et al*., 1990), suggesting that tdOFF responses from the IC likely have a direct excitatory synaptic effect on thalamic neurons, rather than serving to shape ongoing thalamic activity by driving local inhibition. IC projections maintain tonotopy to the MGN, and tdOFF responses in the IC could provide inputs to the thalamus to generate spatially distinct OFF responses in the cortex (Baba *et al*., 2016; Liu *et al*., 2019). However, cortical circuits can also generate OFF responses in the AC (Sollini *et al*., 2018; Bondanelli *et al*., 2021; Solyga & Barkat, 2021), making it difficult to predict how IC tdOFF responses specifically influence downstream cortical processing. The IC also sends GABAergic projections to the MGN (Ito *et al*., 2009; Beebe *et al*., 2018), and IC GABAergic neurons encode sound offset (Wong & Borst, 2019), but whether GABAergic OFF responding neurons are thalamic-projecting or contain tdOFF patterns is currently unknown, but could provide additional tuning of thalamic OFF responses before ascending to the AC.

We previously found that under similar experimental conditions, tone-evoked suppression and tdOFF responses were not present in the IC or the AC in young mice (up to P21) (Babola *et al*., 2018; Kellner *et al*., 2021; Kersbergen *et al*., 2022), suggesting that both are developmentally regulated and arise with the maturation of the auditory system after ear canal opening (~P12 – P14). For mouse AC neurons containing both ON and OFF responses, their receptive fields initially overlap (up to age P23), but diverge by adulthood (> P60), which aids in direction selectivity (Sollini *et al*., 2018). Age-dependent cortical ON/OFF asymmetry is consistent with the developmental emergence of tdOFF responses we see here in the IC (> P56). Thus, in addition to providing ascending OFF responses, the appearance of tdOFF responses from the IC could contribute to AC development.

Sound OFF responses are important for gap detection, the perception of sound duration and termination (Li *et al*., 2021; Solyga & Barkat, 2021), temporal integration (Song *et al*., 2024), and shaping sweep direction selectivity (Sollini *et al*., 2018). What is the perceptual significance of maintaining tonotopically distinct ON and OFF pathways in the auditory system? We found that tdOFF responses resided in a lower frequency space than ON responses in the IC, consistent with previous reports that the characteristic OFF frequency is higher than ON in some IC neurons (Heeringa & van Dijk, 2014; Wong & Borst, 2019). (Baba *et al*., 2016; Liu *et al*., 2019), but, to our knowledge, little work has been done to understand whether OFF tonotopy could be perceived as a separate sound frequency or simply improves the perception of sound offset. Opto-or chemogenetically inhibiting tdOFF responses while maintaining ON and sustained responses in the IC could provide an experimental means to determine their influence on cortical OFF patterns, influence on cortical maturation, and their behavioral role in sound perception.

### Hearing loss preferentially diminishes sideband suppression and tdOFF responses

Hearing loss increases the sound intensity threshold required to evoke a neural response, but also triggers compensatory central gain enhancement in centers throughout the brainstem, thalamus, and cortex that process sound information (Auerbach *et al*., 2014; Palmer & Berger, 2018). A hallmark of gain enhancement is that suprathreshold sound intensities evoke neural responses of similar or potentiated magnitude compared to normal hearing (i.e., similar number of action potentials, field potential amplitude, or fluorescence amplitude), increasing the slope of sound-evoked rate level functions (Auerbach *et al*., 2014; Asokan *et al*., 2018).

Following loud noise exposure as a model of hearing loss, mice exhibited elevated thresholds and increased rate level functions using widefield imaging in the IC. In addition, the temporal waveform of neural responses changed after noise exposure: sideband suppression and tdOFF responses were abolished, and neural activity was less adaptive during the tone. As a result, the spatiotemporal landscape of sound processing was profoundly altered after hearing loss, even though ON response amplitudes were comparable, suggesting similar recruitment of local IC circuits to normal hearing conditions.

Decreased synaptic inhibition and its influence on sound processing is thought to be a core mechanism of central gain enhancement after hearing loss (Auerbach *et al*., 2014; Palmer & Berger, 2018). The loss of sideband suppression in the IC after loud noise exposure supports this hypothesis. The correlated loss of tdOFF responses provides further support for the link between evoked inhibition and tdOFF responses (see above) and provides a potential mechanism for the loss of tdOFF responses after loud noise exposure. Thus, central gain enhancement appears to prioritize ON responses and neural activity during the presence of sound, though this may come at the expense of other sound processing features, such as adaptation, suppression of sideband activity, and tdOFF responses.

In addition to elevated thresholds, hearing loss and noise trauma impair gap processing, as well as speech and sound discrimination in noise, which are attributed to deficits in central sound processing (Caspary & Llano, 2018). Experimentally diminishing OFF responses replicates some of these perceptual impairments (Weible *et al*., 2014; Liu *et al*., 2019; Solyga & Barkat, 2021), suggesting that weakened OFF responses, including tdOFF, with hearing loss may contribute to known hearing deficits beyond sound detection (Kopp-Scheinpflug *et al*., 2018). We used a noise exposure protocol that induced a large shift in sound intensity thresholds (see Methods), though milder protocols induce temporary shifts, allowing thresholds to return to normal hearing levels over time (Kujawa & Liberman, 2009). It is not yet known if IC sideband suppression and tdOFF responses return to normal with hearing thresholds, and whether these changes correlate with improved gap detection and sound discrimination. Addressing these questions will inform whether tdOFF responses are needed to improve sound processing following hearing loss, or whether tdOFF might serve as a therapeutic biomarker for restoring hearing.

## Acknowledgements

Thank you to Dr. Kali Burke for guidance and technical assistance with the loud noise exposure experiments, and Dr. Calvin Kersbergen for technical assistance with sound stimulation and ABR recordings. Thank you to the members of the Bergles Lab for their invaluable input.

## Additional Information

### Data availability

All data reported in this paper and the original code used for analysis will be shared by the lead contact, Dwight E. Bergles, upon request.

